# In vitro implementation of robust gene regulation in a synthetic biomolecular integral controller

**DOI:** 10.1101/525279

**Authors:** Deepak K. Agrawal, Ryan Marshall, Vincent Noireaux, Eduardo D Sontag

## Abstract

Feedback mechanisms play a critical role in the maintenance of cell homeostasis in the presence of disturbances and uncertainties. Motivated by the need to tune the dynamics and improve the robustness of synthetic gene circuits, biological engineers have proposed various designs that mimic natural molecular feedback control mechanisms. However, practical and predictable implementations have proved challenging because of the complexity of synthesis and analysis of complex biomolecular networks. Here, we analyze and experimentally validate a first synthetic biomolecular controller executed *in vitro.* The controller is based on the interaction between a sigma and an anti-sigma factor, which ensures that gene expression tracks an externally imposed reference level, and achieves this goal even in the presence of disturbances. Our design relies upon an analog of the well-known principle of integral feedback in control theory. We implement the controller in an *Escherichia coli* cell-free transcription-translation (TXTL) system, a platform that allows rapid prototyping and implementation. Modeling and theory guide experimental implementation of the controller with well-defined operational predictability.

## INTRODUCTION

Robustness against perturbations and uncertainties is fundamental to biological systems that continuously sense and respond to their environment. At the cellular level, it is often desired to maintain precise control over a variety of molecular components and pathways to achieve complex behaviors that require the interaction of intracellular or extracellular biomolecules.^1,2^ This is often achieved by tightly regulating gene expression in such a way that it follows a desired set point independent of exogenous or endogenous disturbances. Feedback is a mechanism that enables organisms to achieve reliable and robust functionality.^3–7^ Feedback mechanisms underlie homeostasis, a phenomenon in which physiological variables are continuously monitored and adjusted so as to maintain a desired equilibrium value which is defined by a set point (also known as a reference signal), in the presence of biological noise that may perturb the natural state of the system.^8–10^

Reference tracking in the presence of disturbances, a classical objective in electrical and mechanical systems, is often solved by incorporating integral feedback to control a process (also known as a plant).^11^ In such a scheme, one wants a variable of interest (called the output) to track a signal (called the reference). This is achieved by incorporating in the controller the mathematical integration of the difference between the reference and the output (the error signal). The “internal model principle” in control theory states, in essence, that the existence of an integrator in the control loop is necessary for the tight regulation of the concentration of an output species in the presence of disturbances or stimuli.^12^ Inspired by these ideas from engineering and control theory, there have been several recent designs^13–16^ and implementations^17–18^ of biomolecular integral feedback controllers. These implementations are not completely satisfactory in that it is very difficult to make quantitative predictive models of their behaviors, in large part due to the biological noise which is found in cellular systems, which makes it hard to achieve precise control over model parameters, thereby complicating the design and feasibility of the system.^19–21^ Plasmid copy number is also limited by the origin of replication, decreasing the adjustability of experiments.^22^

In this work, we exploit the versatility of an all *E. coli* TXTL platform to prototype a biological controller circuit. TXTL reactions contain the native transcription, translation, and metabolic machineries^23,24^ required to achieve gene expression over at least ten hours. As opposed to a living host, in a TXTL reaction one can precisely set the concentrations and stoichiometries of DNA parts, and thus finely tune gene circuits easily. TXTL reactions are typically performed at the microliter scale or above, far from any biological noise, thus allowing flexibility in designing and optimizing the genetic network.^25^ Experimental disturbances, such as those perturbing the amount of DNA or other reaction components, can be carried out at any time. TXTL reactions are executed in high-throughput, facilitating the rapid characterization of dynamic circuits. By virtue of such advantages, several synthetic gene circuits have been implemented in TXTL with a successful modeling framework.^26–29^

In this article, we constructed a synthetic biomolecular integral controller that precisely controls the expression of a target gene. Our strategy relies on a molecular sequestration reaction that enables error computation between the reference and the output signal while exploiting the natural interaction between the *E. coli* sigma factor σ28 and the anti-σ28, FlgM.^30^ We demonstrate that the output is linearly proportional to the input, in other words, it tracks the reference signal, and this happens for a large dynamic range of input only in the closed-loop configuration (when the sequestration reaction is active). We develop an ordinary differential equation (ODE) model and perform systematic TXTL experiments to parameterize and validate the model. We then successfully predict the controller response in various reaction conditions using the parameterized model. When disturbances are added, only the closed-loop controller enables the output to reject the disturbances. Our results demonstrate that our synthetic biomolecular controller is capable of regulating gene expression robustly in an *E. coli* TXTL toolbox. We anticipate that such an approach could be useful for diagnostics applications,^31^ for constructing dynamical systems *in vitro^32^* or for programming synthetic cell systems.^33,34^

## RESULTS

### Designing an integral feedback controller

Our primary goal is to construct a genetic network which can accurately regulate the expression of a target gene in such a manner that the concentration of a desired (“output”) protein follows a “reference” or input signal linearly, for a large dynamic range of input values. The output should remain at the same concentration when an unknown disturbance affects the system. These desired characteristics of the network should be maintained independently of changes in reference or input values.

In electrical and mechanical control systems, “integral feedback” controllers are routinely used in order to achieve reference tracking in the presence of perturbations and uncertainties. Motivated by this analogy, various possible designs of such controllers have been discussed substantially in the context of biological systems.^13–16,26^ The present work is inspired by our previous work,^26^ in which we introduced a computational design based on RNA based controllers; no experimental validation was provided. Here, we start from that design, modifying it to allow for direct genetic regulation and provide an experimental validation. In this approach, the goal of the controller is to ensure that the intermediate output Z follows the reference signal, which is a scaled value of input P_X_ (Fig. 1a). To determine the deviation of the output from the reference signal, a comparison between both is required without affecting their activity. For that, PX and Z are sensed internally using biochemical reactions such that X and Y represent the input signal and the output signal, respectively. When the output is smaller than the reference signal, the error signal X_FREE_ (X-Y) is detected, and this signal is used in order to correct deviations of the output from the reference signal. For a case when the output is larger than the reference signal, free Y sequesters X to reduce the output. One of the key reasons that this controller is able to improve substantially upon that in^26^ is that it implements an effective error computation through protein interactions; in contrast, RNA-based designs suffer from the fact that RNAs degrade much faster than proteins, making an effective error computation very hard to implement experimentally.^35^ In contrast, the degradation of the proteins used in this work can be ignored.

**Figure 1.**
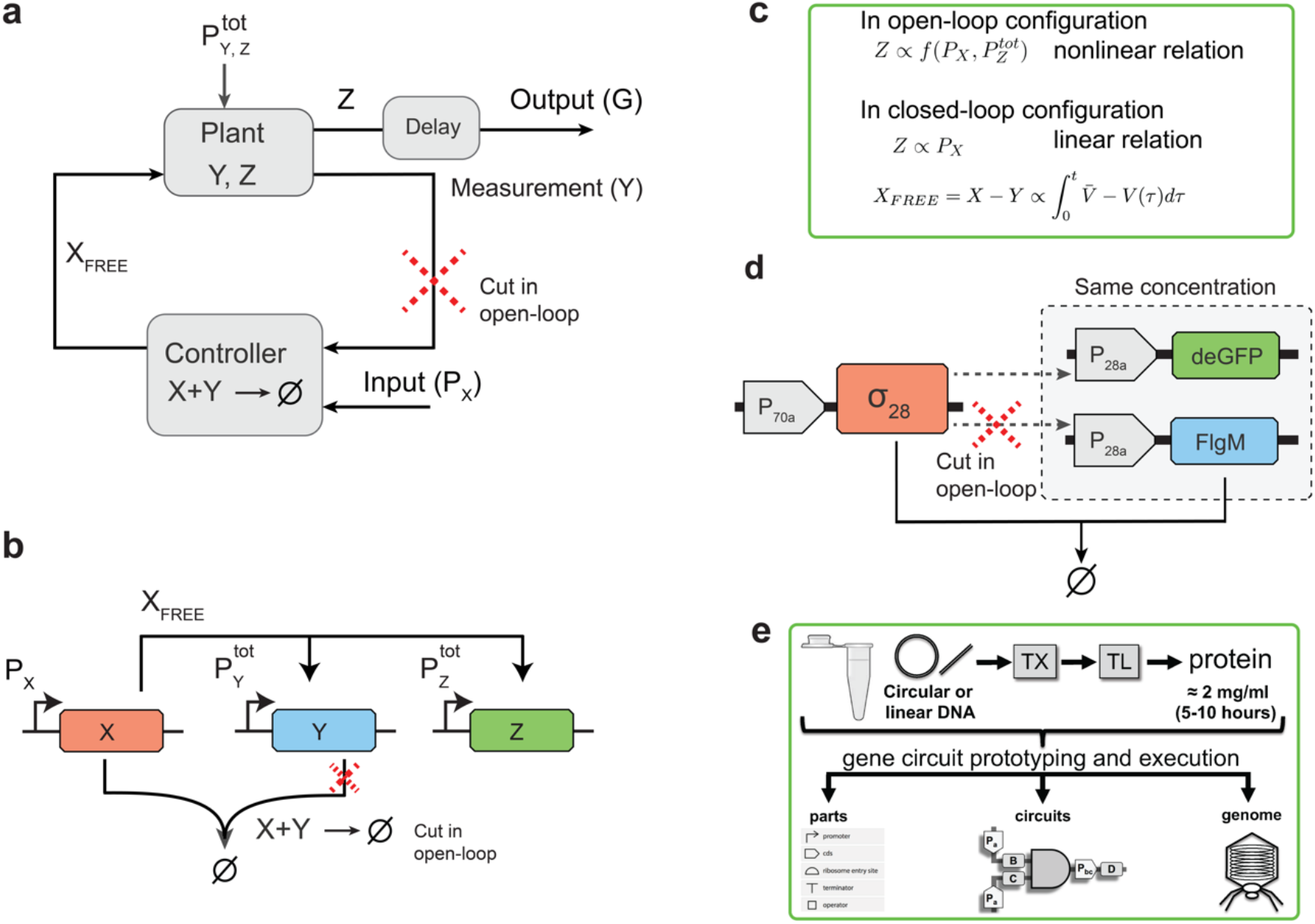
The synthetic biological integral controller. (a) Block diagram of a typical closed-loop controller. (b) Design of the synthetic biological integral controller. The reference is set by the input DNA PX and the intermediate output Z is measured through Y. Error computation is achieved through a molecular sequestration reaction between X (which tracks the reference signal P_X_) and Y (which acts as a proxy for the output Z). Here the red-color cross represents the open-loop configuration of the controller where the feedback signal Y is absent. XFREE is a transcriptional activator that regulates the expression of output Z and Y simultaneously. (c) In the closed-loop configuration, the error signal (XFREE) is the difference between X and Y. To produce an output that is independent of the disturbances in P_Y_^tot^ and P_Z_^tot^, the error signal is mathematically integrated by the controller. Here *<u>V</u>* is the steady state value of V (See SI Note S1). In the absence of the feedback (which is the open-loop configuration) the output depends nonlinearly on P_X_ and P_Z_^tot^ such that any disturbance in P_Z_^tot^ may perturb the output. (d) Experimental implementation of the integral controller. Three plasmids are used, P70a-S28, expressing the *E. coli* sigma factor 28 from a sigma 70 promoter, P28-FlgM, expressing the antisigma 28 (FlgM) from a sigma 28 promoter, and P28a-deGFP, expressing the reporter deGFP from a sigma 28 promoter. In the open-loop controller, instead of FlgM, mSA is expressed, which is not sequestered by sigma 28, nor does it directly interact with any reaction rates. For simplification σ28, FlgM and immature deGFP are denoted as X, Y and Z while promoters P70a and P28a are denoted as P_X_ and P_Z_^tot^ (same as P_Y_^tot^) respectively. (e) Overview of the *E. coli* cell-free toolbox for prototyping and executing parts and circuits *in vitro.*

Our biomolecular implementation of an integral feedback controller requires three genes: an input gene x, under the promoter P_X_, an output gene z, under the promoter P_Z_^tot^ (the target gene) and a proxy gene *y,* under the promoter P_Y_^tot^ for the target gene (Fig. 1b). Genes x, *y* and z encode for respective proteins X, Y, and Z. Note that *x, y* and z have the same concentrations and correspond to the same biochemical species as PX, P_Y_^tot^ and P_Z_^tot^ respectively. The protein X acts as a transcriptional activator for promoters P_Y_^tot^ and P_Z_^tot^. For Y to truly represent Z, it is necessary that the same promoter must be used to express Y and Z, therefore P_Y_^tot^ and P_Z_^tot^ are identical. An error computation is achieved through a molecular sequestration between X and Y such that when X binds to Y or vice versa, both proteins become biologically inactive (a phenomenon, also known as annihilation).^13^ Only free X (X_FREE_, not bound to Y) can activate transcription. It is also required that the synthesis of Y and Z should be overall regulated by XFREE. For that, basal expression from the genes *y* and *z* should be negligible so that in the absence of X_FREE_, the production of Y and Z is negligible. Here and elsewhere, the term “closed-loop configuration” means that the feedback is present through the sequestration reaction; otherwise, when the feedback mechanism is not present, the controller is referred to as in the “open-loop configuration”. We establish that reference tracking is only possible in the closed-loop case, where the output linearly depends on the concentration of P_X_ (Fig. 1c). In the open-loop case, the output depends nonlinearly on the concentration of P_X_ and P_Z_^tot^. Because the error signal (XFREE) is mathematically integrated over time in the closed-loop case, the output should be able to follow the reference signal even when disturbances are added to the plant (Fig. 1a and c).

To test the controller experimentally in TXTL, we employed three plasmids (Fig. 1d): P70a-S28, expressing *E. coli* sigma factor 28 from a sigma 70 promoter; P28a-FlgM, expressing the antisigma 28 (FlgM) from a σ_28_ promoter; and P_28a_-deGFP, expressing the reporter deGFP from a σ_28_ promoter. In the open-loop controller, FlgM is replaced by mSA (same protein size), which is not sequestered by σ_28_, nor does it directly interact with any reaction rates. Here and elsewhere, for simplification, σ_28_, FlgM and immature deGFP are denoted as X, Y and Z, while promoters P_70a_ and P_28a_ are denoted as P_X_ and P_Y_^tot^ (same as P_Z_^tot^) respectively, and the mSA control gene is denoted as *yc* and the respective promoter as P_YC_^tot^.

### The output tracks the input linearly in the closed-loop configuration

Our first goal was to establish that when the controller operates in the closed loop configuration, the output follows linearly the changes in the concentration of input P_X_, thus tracking the reference signal. A nonlinear dependence of the output on P_X_ would suggest otherwise. To test this in TXTL reactions, we added 0.1-0.7 nM P_X_ and 1 nM each of P_Z_^tot^ and either P_YC_^tot^ for the open-loop operation (Fig 2a) or P_Y_^tot^ for the closed-loop operation (Fig 2b). In the open-loop case, we found that changes in the deGFP concentrations depend nonlinearly on the changes in the input concentration of PX (Fig. 2c), suggesting that the output is unable to track the reference signal over the tested range. In contrast, the closed-loop endpoint deGFP concentrations were linearly proportional to the concentration of P_X_ (Fig.2d), suggesting reference tracking.

**Figure 2.**
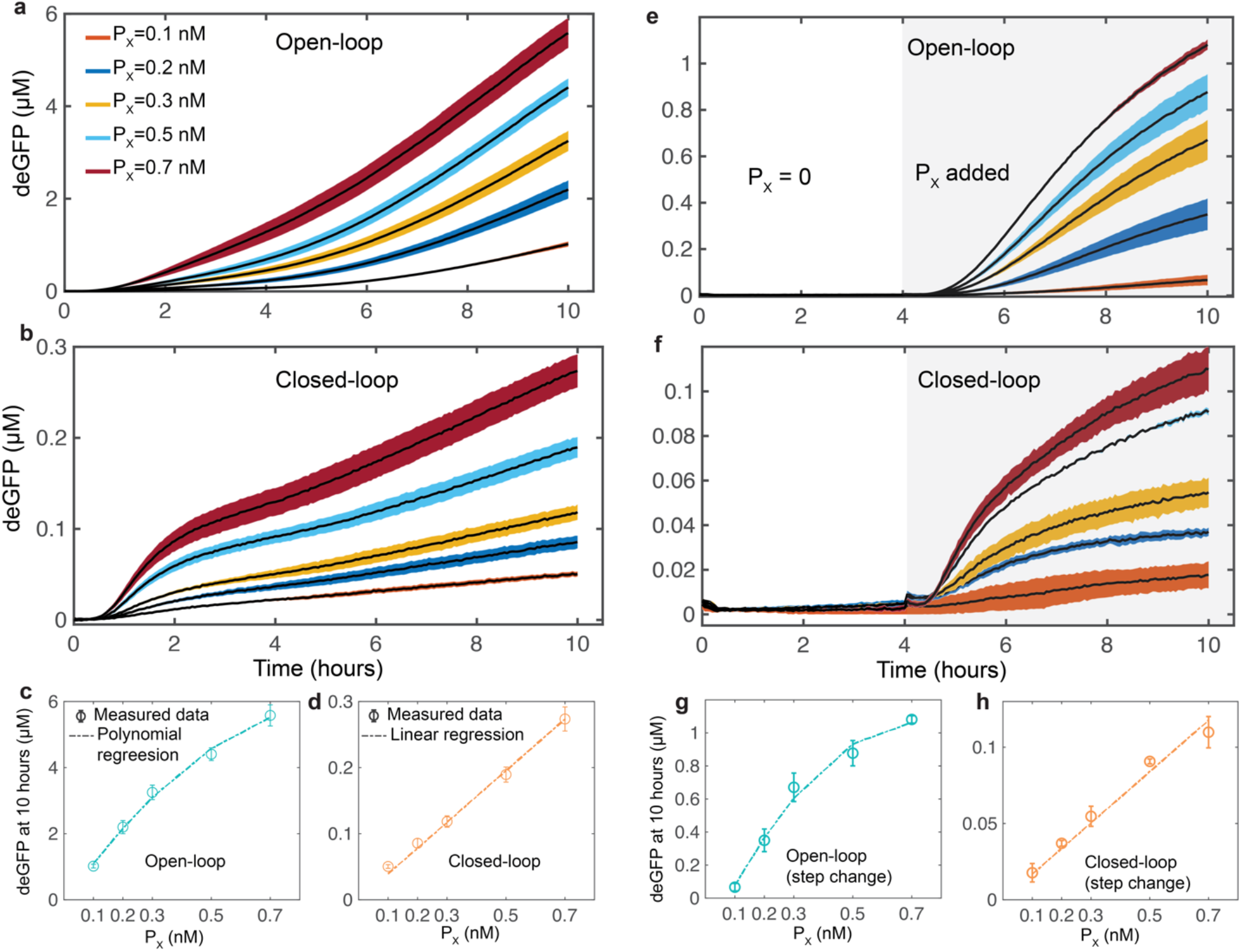
The output tracks the input signal linearly in the closed-loop configuration. (a,b) TXTL measurement of the response of the integral controller in the (a) open-loop and (b) closed-loop configurations at different initial concentrations of P_X_ (0.1 – 0.7 nM) while initial P_Y_^tot^ and P_Z_^tot^ were both 1 nM each. (c,d) Summary of the measured deGFP response of the controller at 10 hours respectively. To disable the feedback in the open-loop case P_Y_^tot^ was replaced by P_YC_^tot^, which expresses a protein that cannot sequester with X. (c-d) Measured response for a step change in P_X_. P_X_ was increased from 0 nM to different concentrations (0.1-0.7 nM) after 4 hours of the reaction in the presence of initial 1 nM of P_Y_^tot^ and P_Z_^tot^ each. (g,h) Summary of the measured deGFP response of the controller at 10 hours respectively. Error are shown in the shaded region and were determined using the standard error of the mean of three or more repeats. A two-degree polynomial was used to fit the measured deGFP endpoints for (c,g) the open-loop case while a linear regression with zero intercept was used to fit the closed-loop response where slope (d) m=0.0108 and (h) m=0.008. A calibration factor was used to convert the measured deGFP fluorescent signal into the concentration.

The output of the closed-loop controller should be able to follow the changes in input signal linearly independent of the time when it is modified. To test this capability, we performed a step change in the input P_X_ concentration during the course of the reaction. For that, we added different amounts of P_X_ to TXTL reactions in the open-loop (Fig. 2e) and closed-loop (Fig. 2f) system after four hours of incubation with 1 nM each of P_Z_^tot^ and either P_YC_^tot^ or P_Y_^tot^, respectively. As mentioned earlier, we observed that the controller’s output follows linearly the input only when operated in the closed-loop configuration (Fig. 2g and h). Note that the deGFP produced in the closed-loop configuration is much smaller than that produced in the open-loop configuration because the activator needed to express the deGFP is sequestered. As a control, we also tested that changes in the concentration of P_YC_^tot^ have no effect on the output (Fig. S1), confirming that the different version of Y that is expressed by *yc* gene does not interact with X. We also found that in the absence of P_X_, deGFP is not produced (Fig. S2), confirming that the production of Y and Z are completely governed by X through the activation reaction.

These experimental observations agree with the expected controller operation. When X is larger than Y, (i.e. the output is lagging behind the reference), X_FREE_ increases the production of Y and Z. As more Y is available in the reaction to sequester with X, X and Y converge to specific values that would allow Z to follow the reference signal. In the absence of the sequestration reaction, error computation is absent (no feedback) and so X directly regulates Z production without comparing with the reference signal.

### Mathematical model and parameterization

To understand the controller operation, we developed a simple coarse-grained model that captures the dynamic response of the controller in both open and closed-loop configurations (Fig. 3a). For that, we consider the synthesis of each protein as a two-step reaction: a transcription reaction for mRNA synthesis and then a translation reaction for the corresponding protein synthesis. The parameters α and β are transcription and translation rates, respectively. Here and elsewhere, subscripts to the parameters indicate the corresponding species. Each mRNA species (U, V and W) has a degradation rate denoted as δ while we ignore the protein degradation rate.^35^ The parameter κ is the sequestration rate. Transcriptional activation is modeled as a one-step reaction, where X binds to the promoters P_Y_ and P_Z_ separately at a rate of ω and dissociates at a rate of v. In the activated state, these genes produce Y and Z proteins at an increased transcription rate, denoted as α^+^. Considering the mass-conservation, we assume that P_Y_^tot^ = P_Y_ + P_Y_^+^ and P_Z_^tot^ = P_Z_ + P_Z_^+^ at all times. An additional reaction is added into the model to account for the maturation of newly synthesized deGFP (Z) into a fluorescent deGFP (G),^24,32^ which is the read-out signal (Fig. 3a). Here onwards, fluorescent deGFP is referred as deGFP. From chemical reactions, we built an ODE model (shown in Fig. 3b) to determine the response of the controller over time.

**Figure 3.**
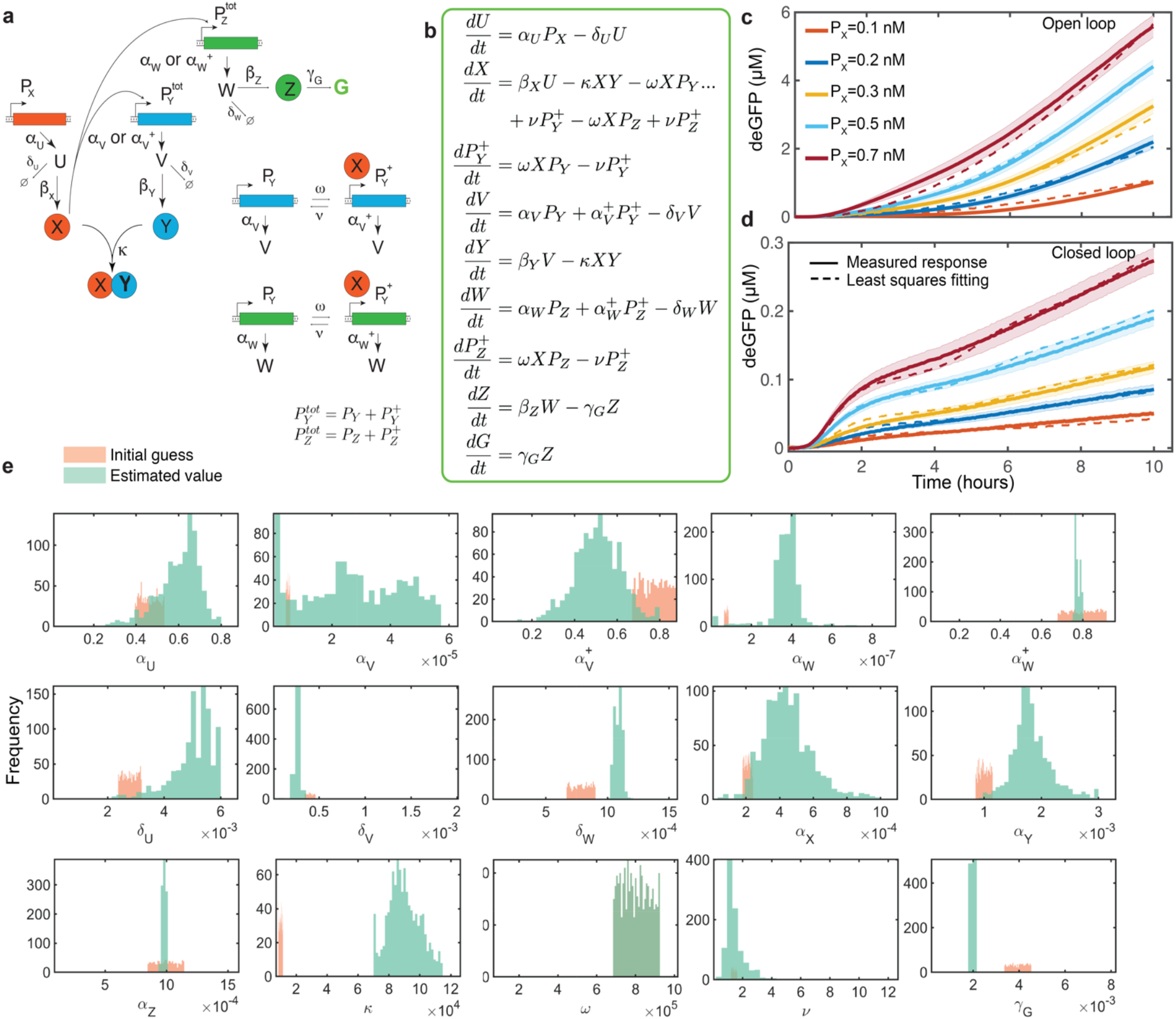
Schematic of the model parametrization process and validation of the model. (a) Detailed reaction network of the controller and the corresponding (b) ODE model. Here U, V and W are the translationally initiated mRNA of the X, Y and Z proteins respectively. The additional reaction was added into the model to consider the maturation of immature deGFP into a fluorescent deGFP (G). Details on how the P_Y_^tot^ and P_Z_^tot^ promoter switch from the inactive (P_Y_, P_Z_) to active (P_Y_^+^, P_Z_^+^) states are shown alongside. (c-d) Comparing the measured deGFP response (solid lines) with the mean of the best-fitted simulation results (dashed lines) for the (c) open-loop and (d) closed-loop cases. ODE model was used to calculate the controller response while initial concentrations of P_Y_^tot^ and P_Z_^tot^ were 1 nM each. Note for the open-loop response αV, αV^+^, δ_V_, α_Y_, κ and P_Y_^tot^ were set to zero. Least squares fitting was used to generate the best-fit model response at different initial guesses of the reaction parameters (see Methods). Experimentally observed error bars are shown in the shaded region while the mean simulated trajectories (dashed line) are shown here within 95% confidence intervals. N=1000. Error bars are from the SEM of at least three repeats. (e) Histograms of the initial guesses and estimated parameters were obtained from 1000 samples that gave the lowest fitting error (see Methods).

An accurate representation of a system model requires determining specific parameter values at which the model quantitatively follows the system dynamics. However, parameter estimation can be nontrivial, as there can be multiple sets of parameter values that may vary by several orders of magnitude and could still fit the measured data. Therefore, to simplify the problem, we isolated the measured responses into two sets, in such a manner that we required fewer parameters to fit a particular set of experimental data. For this, we first found the model parameter values that provided the best fit to the measured open-loop response, since the number of parameters involved in the open loop case are less than in the closed-loop case (see Methods). We then used these parameter values to fit the closed-loop response while allowing only the remaining parameters to vary (see Methods). Moreover, we started the model fitting manually with an initial guess of parameters derived from the literature.^23–29,35^ Once we found the possible values that provide a qualitative agreement between the model and the measured response, we used an iterative least-squares fitting procedure to find a range of parameter values that gave us the best fit (see Methods). The means of the resulting parameters (Table 1) were then used along with the ODE model (Fig. 3b) to calculate the mean trajectories with 95% confidence intervals (Fig. 3c and d). Histograms of the input and estimated parameter distributions are shown in Fig. 3e.

**Table 1:** Estimated model parameters obtained from the least squares fitting for the ODE model shown in Fig. 3b. Note for the open-loop response α_V_, α_V_^+^, δ_V_, α_Y_, κ and P_Y_^tot^ were set to zero. The error values were determined using the standard error of the mean. N=1000.

| Parameters | Values | Error | Units |
| --- | --- | --- | --- |
| $\alpha_U$ | 0.60 ± | 0.0031 | $s^{-1}$ |
| $\alpha_V$ | $2.75 \times 10^{-5}$ ± | $5.19 \times 10^{-7}$ | $s^{-1}$ |
| $\alpha_V^+$ | 0.51 ± | 0.0034 | $s^{-1}$ |
| $\alpha_W$ | $3.68 \times 10^{-7}$ ± | $2.90 \times 10^{-9}$ | $s^{-1}$ |
| $\alpha_W^+$ | 0.78 ± | 0.0005 | $s^{-1}$ |
| $\delta_U$ | 0.00501 ± | $2.43 \times 10^{-5}$ | $s^{-1}$ |
| $\delta_V$ | 0.00026 ± | $2.00 \times 10^{-6}$ | $s^{-1}$ |
| $\delta_W$ | 0.00109 ± | $1.23 \times 10^{-6}$ | $s^{-1}$ |
| $\beta_X$ | 0.00045 ± | $4.64 \times 10^{-6}$ | $s^{-1}$ |
| $\beta_Y$ | 0.00183 ± | $1.10 \times 10^{-5}$ | $s^{-1}$ |
| $\beta_Z$ | 0.00098 ± | $8.94 \times 10^{-7}$ | $s^{-1}$ |
| $\kappa$ | $8.99 \times 10^{+4}$ ± | $3.22 \times 10^{+2}$ | $M^{-1} s^{-1}$ |
| $\omega$ | $8.02 \times 10^{+5}$ ± | $2.19 \times 10^{+3}$ | $M^{-1} s^{-1}$ |
| $\nu$ | 1.54 ± | 0.03 | $s^{-1}$ |
| $\Upsilon_G$ | $1.95 \times 10^{-3}$ ± | $7.88 \times 10^{-6}$ | $s^{-1}$ |

To cross-validate the parameterized model, we predicted the deGFP response for two different settings of input conditions. In the first setting, the concentration of PX was increased from 0 nM to 0.1-0.7 nM (in a step manner) after 2 hours of incubation in the presence of initial 0.7 nM of P_Y_^tot^ and P_Z_^tot^ each (Fig. 4a, 4b and Fig. S3). In the second setting, the concentration of P_Y_ was varied from 0.2 nM to 1 nM at the beginning of the reaction while keeping the initial concentration of P_X_ and P_Z_ at 0.7 nM each (Fig. S4). In all the cases, the predicted responses followed the measured responses closely.

**Figure 4.**
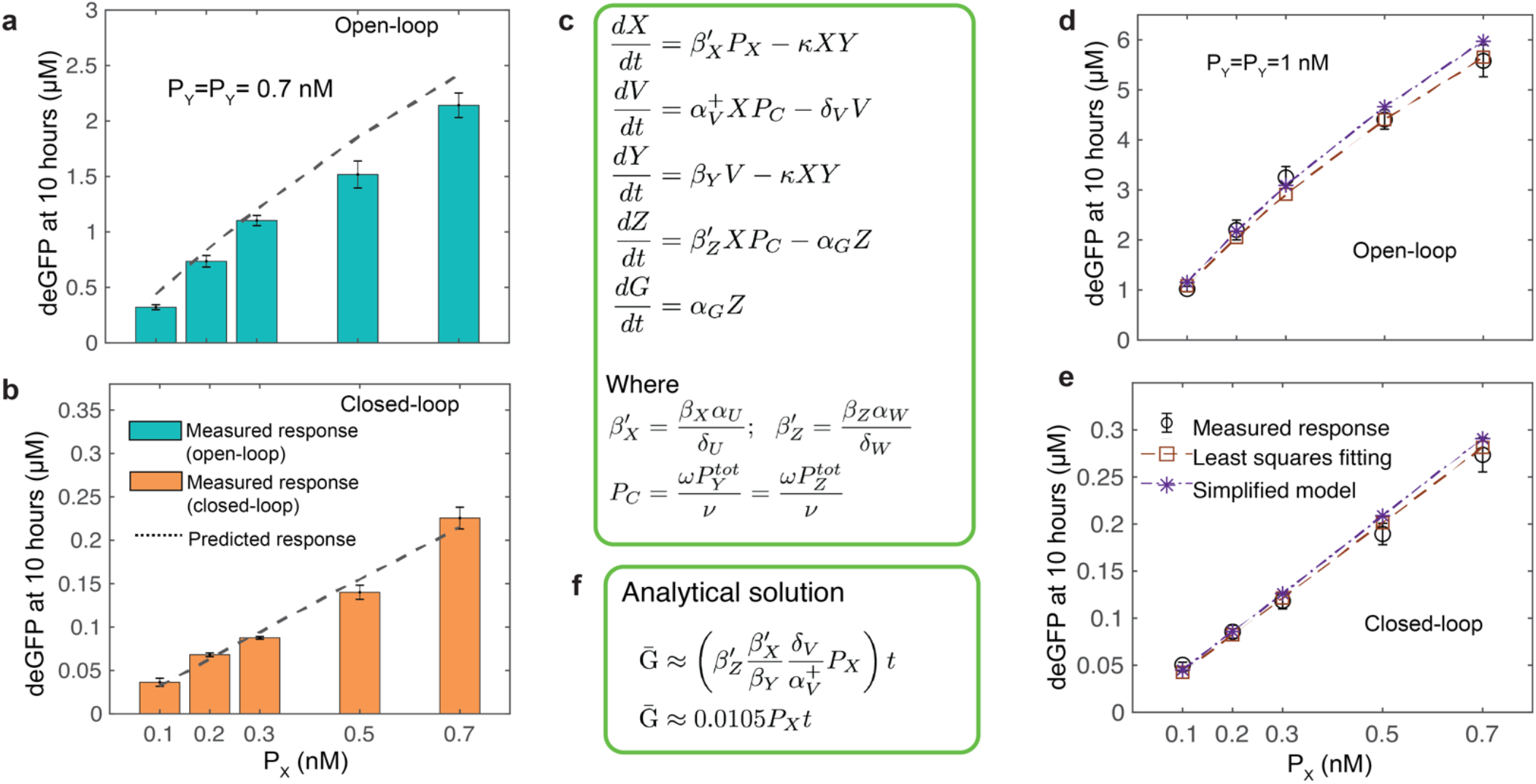
Model simplification and predicting the controller response. (a-b) Predicting the (a) open-loop and (b) closed-loop controller response for a step change in P_X_. P_X_ was increased from 0 nM to different concentrations (0.1-0.7 nM) after 2 hours of incubation in the presence of initial 0.7 nM of P_Y_^tot^ and P_Z_^tot^ each. The ODE model shown in Fig. 3b was used to determine the response with parameters shown in Table 1. (c) Simplified ODE model of the controller. (d) Comparing the simplified model response with the original model response for the (d) open and (e) closed-loop cases. (d) Analytical solution of the reporter protein deGFP over time that is defined as a reference signal. Using the parameters values shown in Table 1, constant rate (shown inside the brackets) was determined.

### Model simplification

To get a better insight into the controller response, further simplification of the proposed model is required. Based on the extracted parameter values, we sought to minimize the number of variables in the model while still capturing the essential dynamics of the controller. From the extracted parameter values, we observed that the basal expression of *y* and *z* genes is almost negligible; therefore we set αv and αw to zero in the simplified model. As the transcriptional activation reaction is much faster than the other reactions involved in the reaction network, we used a quasi-steady state approximation to replace the ODEs of P_Y_^+^ and P_Z_^+^ by their steady-state expressions (see SI Note 1). As the transcription reaction is much faster than the translation reaction, a similar approximation was used to model the synthesis of X and Z using single reactions for each, while keeping the two-step synthesis of Y to ensure that we consider an appropriate delay in the overall system dynamics. This leads to a simplified model (shown in Fig 4c), which can produce dynamic response similar to that of the original model (Fig. 4d, e, and Fig. S5).

We can now use the simplified model to determine analytically how the output depends on the reaction parameters and the input. In our implementation of the controller, the reporter protein deGFP (G) has no degradation tag, consequently we cannot observe a steady state behavior in the measured response. However, a linear dependence of the time derivative of G on Z (see Fig. 4c), suggests that when Z reaches a steady-state, G should increase at a constant rate over time (see Fig. S6). To determine this constant rate, we first used the simplified model to determine the steady state value of Z (see SI Note S1) and then we calculated the analytical solution of G over time, as shown in Fig. 4f. The output G increases at a constant rate that is the scaled value of the input P_X_ and used as a reference signal.

Note that because of the quasi-steady state approximation (see SI Note S1), the analytical solution can only be used to determine the closed-loop controller response when all other reactions reached a steady-state. We then used the extracted parameters to calculate this constant rate and found that it closely matches the observed response shown in Fig. 2d, confirming that the controller output tracks the reference signal in the closed-loop configuration. Further insight into the controller operation can be gained by analyzing the simplified model to determine how the error signal is processed in the closed loop configuration of the controller. The analytical equation of XFREE clearly shows that the error signal (which is X-Y) is integrated mathematically by the controller (see SI Note S1).

### Closed loop control enables disturbance rejection

One of the main theoretical advantages of integral feedback controllers is their ability to minimize the effect on the output of disturbances on the system.^9^ This is due to the effective error computation with an integral operation that allows the controller to maintain the desired output even when a disturbance affects the system. In the aforementioned section, we showed that the implemented controller can be interpreted as an integral feedback mechanism (see SI Note S1). Therefore, we expect that the closed-loop system is able to reject disturbances in an appropriate sense. To test this, we introduced disturbances in the concentration of the biochemical species P_Y_^tot^ and P_Z_^tot^. In practical settings, variation in the DNA concentration is one of the most biologically relevant parameters. This is because *in vivo* gene concentration can vary significantly due to fluctuations in plasmid copy number,^36^ and several designs have been proposed in order to ameliorate the effect of copy number variation.^37,38^ Notably, the TXTL reaction platform allows us to design such an experiment, where DNA template concentrations can be changed at any time during the course of a reaction due to the TXTL reaction settings.

In this design, the reference signal is independent of the amount of *y* and *z* genes (in Fig. 4f) so that any disturbances in these species should not perturb the deGFP response when the controller is operating in the closed-loop configuration. To test this, we first used the ODE model (Fig. 3b) to predict the controller response in the open-loop and closed-loop configurations where the concentrations of P_Y_^tot^ and P_Z_^tot^ were varied from 0.2 to 0.7 nM, keeping a fixed 0.2 nM initial concentration of P_X_. We observed a less than 10% variation in the deGFP response for the closed-loop case, compared to over 300% variation in the open-loop case (Fig. S7). In the closed-loop operation, an increase in P_Y_^tot^ increases the amount of Y, but as more Y is available to sequester X, less X_FREE_ is available to activate the production of Y and Z. Even though P_Z_^tot^ was increased, X_FREE_ is reduced such that the reporter protein (G) remains the same, independently of the amount of P_Z_^tot^ (Fig. S8). For the open loop case, as there is no feedback, an increase in P_Z_^tot^ significantly increases the production of G (Fig. S9).

Encouraged by these results, we experimentally tested the same conditions in TXTL reactions. We added 0.2 nM P_X_ and increasing concentrations of P_Y_^tot^ = P_Z_^tot^, from 0.2 nM to 0.7 nM to reactions, and tracked the deGFP output over time for the open-loop (Fig 5a) and the closed-loop (Fig 5b) configurations. In the open-loop case, the output signal increased with the increasing concentration of P_YC_^tot^ = P_Z_^tot^ due to the lack of feedback (Fig 5c). However, in the closed-loop case, the output signal was independent of the increasing P_Y_^tot^ = P_Z_^tot^, thus confirming that the controller rejected the disturbance in their concentrations (Fig 5d), as predicted by integral feedback theory

**Figure 5.**
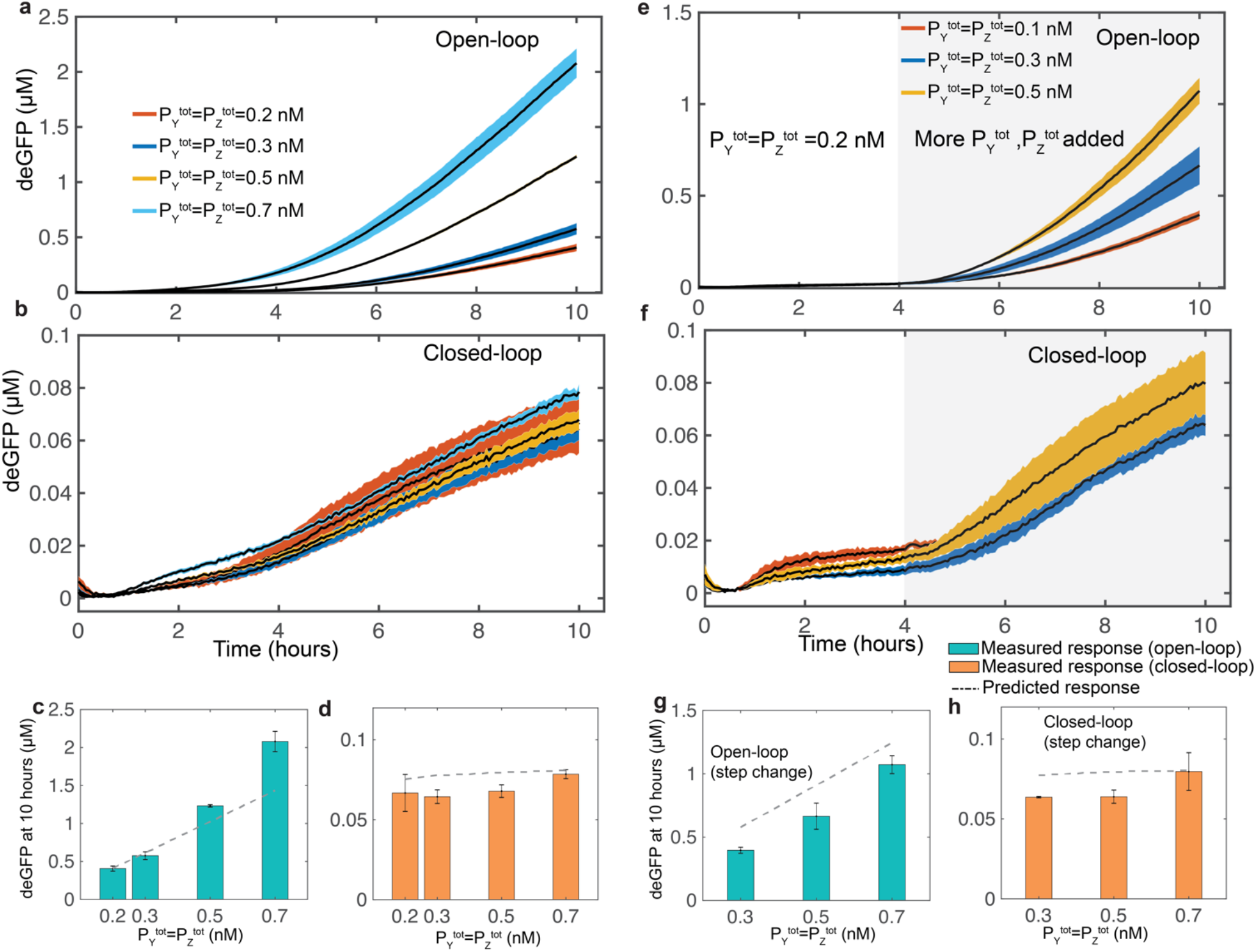
Closed-loop operation enables the controller to achieve robustness to disturbances on the system. (a,b) Measured deGFP response of the controller in the presence of disturbances in the concentration of P_Y_^tot^ and P_Z_^tot^ (0.2 – 0.7 nM) for the (a) open-loop and (b) closed-loop cases while initial P_X_ was 0.2 nM. (c,d) Summary of the measured deGFP response of the controller at 10 hours respectively. To disable the feedback in the open-loop case P_Y_^tot^ was replaced by P_YC_^tot^, which expresses a protein that cannot sequester with X. (e,f) Measured response of the controller when the disturbance in P_Y_^tot^ and P_Z_^tot^ was added in a step manner. Additional P_Y_^tot^ and P_Z_^tot^ were added (0.1-0.5 nM) after 4 hours of the reaction in the presence of initial 0.2 nM of P_X_, P_Y_^tot^ and P_Z_^tot^ each. (g,h) Summary of the measured deGFP response of the controller at 10 hours respectively. The error bars are shown in the shaded region and were determined using the standard error of the mean of three or more repeats. The predicted response for each case was determined using the ODE model shown in Fig. 3b with parameters shown in Table 1.

Similarly to reference tracking, rejection of disturbances should be independent of the way that the disturbance is introduced in the system. To test this, we characterized the controller response to a step change in the concentrations of P_Y_^tot^ and P_Z_^tot^ as disturbances. We started the reaction with P_X_, P_YC_^tot^ and P_Z_^tot^ concentrations each set to 0.2 nM, and after four hours of incubation, additional P_YC_^tot^ and P_Z_^tot^ were added (see Methods) (Fig. 5e). For the closed-loop case, instead of P_YC_^tot^, P_Y_^tot^ was added in the same amounts (Fig. 5f). The output was not perturbed in the presence of the disturbances (Fig. 5g,h) only in the closed-loop case, as expected based on our understanding of the controller. We also tested the controller’s response for a wide range of disturbances when different initial concentrations of DNA were used, and the disturbance was added in different amounts (See Fig. S10, S11). In each case, we found a similar robust controller response in the closed-loop settings.

## DISCUSSION

In this work, we constructed a synthetic biomolecular integral controller circuit for effective and robust control of gene expression. We demonstrated that in the closed-loop configuration, the output followed the input signal linearly over a wide range of conditions, even under step changes in input. By harnessing the natural interaction between the sigma factor σ28 and the anti-σ28, FlgM, a strong sequestration reaction is realized that allowed an effective computation of the error between the reference and the output signal. This error signal is then integrated mathematically. Because of this, and as predicted by theory, the closed-loop controller output rejects the disturbances introduced in the DNA concentration. In contrast, the open-loop controller is unable to reject the disturbances, as noticed by the large variations of the output signal. This illustrates the advantage of closed-loop architectures, where an error computation is employed.

Mathematical models play an important role in understanding complex synthetic networks. They provide insight into the operation of networks, and serve to guide experimentation. Here, we developed an ODE model for the integral controller that quantitatively explains the transient as well as the steady-state behavior of all the species involved in the system. We were able to obtain an effective model parameterization, by isolating parameters for each set of experimental data before finding their optimum values to achieve the best fit. Even though the presented model is a coarse-grained mechanistic model, it enabled us to explain the measured response of the controller, and can also predict dynamic trajectories for a wide range of operating conditions accurately. Based on the extracted parameter values, we derived a simplified version of the original model that is as effective as the original model and can be used for further theoretical analysis in order to gain deeper insight into the operation of the controller.

We have limited the model to the measured dynamic trajectories for up to 10 hours. Experimentally, after 10 hours, we observed effects of resource limitations (Fig. S12). We assume that the TXTL reactions have an unlimited source of energy for the first 10 hours in the range of plasmid concentrations used. After this point, we start to see a decrease of transcription and translation rates consistent with the depletion of energy resources and biochemical changes such as pH drop.^24^ The current ODE model assumes an unlimited energy source and therefore ignores the experimental results when that assumption no longer holds. Future work could incorporate resource competition and depletion. Because the controller was implemented in a TXTL reaction platform at the scale of a few microliters, a deterministic model was used while ignoring the biological noise. For *in vivo* applications, it may be desirable to extend the deterministic model to a stochastic model to consider intrinsic the biological noise.^36^ We used a molecular sequestration reaction to realize the error computation. This strategy has been used previously to design closed-loop biomolecular controllers for reference tracking and disturbance rejection.^13–18^ Although our design is a variation of those in,^13,26^ it differs from them in substantial ways in its biological instantiation. Our results are unique in the sense that we have a well-characterized controller that is realized in all *E. coli* TXTL system, where biological noise is negligible compared to *in vivo* implementations. In the latter, biological noise plays a dominant role in governing the system dynamics.^18^ Because of this, we can not only precisely regulate gene expression, but can also accurately predict the dynamic response of the integral controller. Our controller design, which uses a genetic network, can also be easily tailored to regulate any other gene of interest such as those involved in controlling metabolic rates, a task which might not be feasible using a post-transcriptional based controller.^13^ We also demonstrate disturbance rejection capabilities of our controller when a constant or a step disturbance is added to DNA concentration (P_Y_^tot^ and P_Z_^tot^); disturbances in DNA concentration are realistic because they typically arise in *in vivo or in vitro* genetic networks. Such experiments are not feasible using an *in vivo* reaction platform, thereby limiting the usage of the controller in rapid prototyping and implementation.

Notably, for the current architecture, disturbance rejection is only possible when perturbations do not directly influence the parameters involved in governing the steady-state response of the reporter protein (shown in Fig. 4f). It is also important to note that the closed-loop controller can only reject the disturbance provided that the genes *y* and z are at the same concentration and that the added disturbance to both is the same, ensuring that at any time during the reaction it holds that P_Y_^tot^ = P_Z_^tot^. To achieve this operating condition *in vivo,* Y and Z could be expressed on the same operon, controlled by a single promoter. Finally, in this work, we demonstrated that the closed-loop controller can robustly control single gene expression of the deGFP fluorescent reporter protein taken as a model process to be controlled. However, we anticipate that the controller design could be extended to tightly regulate multiple genes that encode other biologically relevant proteins simultaneously or could be employed within a complex network system where multiple processes required tight regulation to improve robustness and performance of the network.

Molecular controllers capable of robust gene regulation are needed in synthetic biology in order to implement more complex circuit networks. The well-characterized and rationally implemented synthetic integral feedback controller we presented here is capable of addressing these challenges to advance biological engineering, and could lead to the development of powerful, synthetic network systems capable of achieving complexity similar to that found at the cellular level, to develop cell-free applications such as calibrated biomanufacturing or programming synthetic cells for specific tasks.

## METHODS

### Mathematical modeling and parameter estimation

The simulated response of the controller was determined by numerically integrating ODE models (Fig. 3b) using MATLAB ode23s solver unless otherwise specified. Initial conditions for each molecular species are described in the figure captions, and the values of reaction parameters are shown in Table 1. For the cases where there is a step change in the DNA concentration over the course of the reaction, similar settings were used to determine the model response numerically.

For parameter estimation, first we found initial guesses of parameter values that qualitatively agreed with the measured open-loop response. We then randomly sampled a set of input parameters from a uniform distribution within a bounded interval (upper and lower bounds of 15% each) centered around the initial guess values. This input set of parameters was then optimized to minimize the error between the model and measured open-loop responses for all five trajectories (shown in Fig. 3c). To find the best fit, the least squares error between the model and the measured response was minimized using the MATLAB fmincon function. During the fitting, each input parameter was allowed to vary from 0.1 to 10 times with respect to the input value. Further constraints were placed on some parameters so that they lie within a feasible range. For example, the activated transcriptional rate must be several orders of magnitude larger than the basal expression (α_V_^+^>>α_V_ and α_Z_^+^>>α_Z_) and the transcriptional rates of the *x* gene and activated *y* and *z* genes should be in the same order (α_X_≈α_Y_^+^≈α_Z_^+^) (See SI Table S1). Because in the open-loop case, a modified version of *y* gene (denoted as *yc*) is expressed from the promoter P_Y_^tot^ and cannot sequester with X, parameters α_V_, α_V_^+^, δ_V_, αγ, κ and P_Y_^tot^ were set to zero. Once we found the optimum set of parameter values that provided the best fit for the open-loop response, these parameter values were then fixed during fitting all five trajectories of the closed-loop response (shown in Fig. 3d) while varying only α_V_, α_V_^+^, δ_V_, α_Y_ and κ. This resulted in a set of 15 parameters that fit both open and closed-loop responses (Fig. 3c and d, respectively). The fitting process was repeated 1000 times, which gave a range for the 16 parameters (Fig. 3e) with 95% confidence interval.

### TXTL Preparation and Reactions

The all *E. coli* cell-free TXTL extract was prepared from BL21 Rosetta 2 from Novagen, as described previously.^23,24^ The TXTL system is commercially available as the product “myTXTL” from Arbor Biosciences. TXTL reactions are composed of 1/3 volume cell lysate, with the remaining 2/3 volume containing plasmids, amino acids, and reaction buffers. All TXTL reactions contained 50 mM HEPES pH 8, 1.5 mM ATP and GTP, 0.9 mM CTP and UTP, 0.2 mg/mL tRNA, 0.26 mM coenzyme A, 0.33 mM NAD, 0.75 mM cAMP, 0.068 mM folinic acid, 1 mM spermidine, 30 mM 3-PGA, 1.5% PEG8000, 30 mM maltodextrin, 3 mM each of 20 amino acids, 90 mM K-glutamate, and 4 mM Mg-glutamate. TXTL reactions were assembled using the Labcyte Echo 550 liquid handler, to volumes of 2 μl in a 96-well V-bottom plate (Corning Costar 3357 with caps Costar 3080) and incubated at 29°C.

### DNA

Plasmids were constructed using standard cloning techniques. Each plasmid contains the untranslated region UTR1, and either the σ70 promoter P70a or the σ28 promoter P28a, all described previously.^23,24,35^ For experiments with step changes in the concentration of DNA, the TXTL reactions were assembled in the same manner, using the Labcyte Echo 550, and incubated in a plate reader at 29°C. Reactions were then taken out of the plate reader, and the additional DNA was added to the reaction using the Labcyte Echo 550. The well plate was then immediately returned to the plate reader. The total time that the well plate was out of the reader and at room temperature was less than two minutes. The step-change of DNA added to the reactions diluted the TXTL reaction by less than 5%. Plasmid sequences can be found in the Supplementary Note S2.

### TXTL Time-Course Fluorescence Measurements

Fluorescence kinetics were performed using the reporter protein deGFP, a truncated version of eGFP that is more translatable in the TXTL system (25.4 kDa, 1 mg/mL = 39.38 μM).^23^ Measurements were carried out on Synergy H1 and Neo2 (Biotek Instruments) plate readers, using an excitation of 485 nm and emission of 525 nm, measuring every 3 minutes. To quantify the concentration of deGFP on the plate readers, a standard curve of intensity vs. deGFP concentration was made using recombinant eGFP (Cell Biolabs Inc.).^35^ All reactions were performed in at least triplicate.

## ACKNOWLEDGMENTS

This material is based upon work supported by the Defense Advanced Research Projects Agency (contracts FA8650-18-1-7800 and HR0011-16-C-01-34), the National Science Foundation (contract 1817936), and the Air Force Office of Scientific Research (contract FA9550-14-1-0060)

## REFERENCES

1 Alon, U., Surette, M. G., Barkai, N. & Leibler, S. Robustness in bacterial chemotaxis. Nature 397, 168–171 (1999).

2 Del Vecchio, D. & Murray, R. M. Biomolecular feedback systems. (Princeton University Press Princeton, NJ, 2015).

3 Dunlop, M. J., Keasling, J. D. & Mukhopadhyay, A. A model for improving microbial biofuel production using a synthetic feedback loop. Syst Synth Biol 4, 95–104, doi:10.1007/s11693-010-9052-5 (2010).

4 Samaniego, C. C., Giordano, G., Kim, J., Blanchini, F. & Franco, E. Molecular Titration Promotes Oscillations and Bistability in Minimal Network Models with Monomeric Regulators. Acs Synthetic Biology 5, 321–333, doi:10.1021/acssynbio.5b00176 (2016).

5 Yi, T. M., Huang, Y., Simon, M. I. & Doyle, J. Robust perfect adaptation in bacterial chemotaxis through integral feedback control. P Natl Acad Sci USA 97, 4649–4653, doi:DOI 10.1073/pnas.97.9.4649 (2000).

6 Barkai, N. & Leibler, S. Robustness in simple biochemical networks. Nature 387, 913–917, doi:10.1038/43199 (1997).

7 Tang, C., Ma, W. Z., Trusina, A., El-Samad, H. & Lim, W. Defining network topologies that can achieve biochemical adaptation. Faseb J 24(2010).

8 Darrasse-Jeze, G. et al. Feedback control of regulatory T cell homeostasis by dendritic cells in vivo. J Exp Med 206, 1853–1862, doi:10.1084/jem.20090746 (2009).

9 Aström, K. J. & Murray, R. M. Feedback systems: an introduction for scientists and engineers. (Princeton university press, 2010).

10 Angeli, D., Ferrell, J. E., Jr. & Sontag, E. D. Detection of multistability, bifurcations, and hysteresis in a large class of biological positive-feedback systems. Proc Natl Acad Sci U S A 101, 1822–1827, doi:10.1073/pnas.0308265100 (2004).

11 Sontag, E. D. Mathematical control theory: deterministic finite dimensional systems. Vol. 6 (Springer Science & Business Media, 2013).

12 J. Huang, A. Isidori, L. Marconi, M. Mischiati, E. D. Sontag, and W. M. Wonham, Internal models in control, biology and neuroscience, in Proc. IEEE Conf. Decision and Control, Miami, Dec. 2018, IEEE Publications, Piscataway, NJ, 2018

13 Briat, C., Gupta, A. & Khammash, M. Antithetic Integral Feedback Ensures Robust Perfect Adaptation in Noisy Biomolecular Networks (vol 2, pg 15, 2016). Cell Syst 2, 133–133., doi:10.1016/j.cels.2016.02.010 (2016).

14 Briat, C., Zechner, C. & Khammash, M. Design of a Synthetic Integral Feedback Circuit: Dynamic Analysis and DNA Implementation. ACS Synth Biol 5, 1108–1116, doi:10.1021/acssynbio.6b00014 (2016).

15 Oishi, K. & Klavins, E. Biomolecular implementation of linear I/O systems. IET Syst Biol 5, 252–260, doi:10.1049/iet-syb.2010.0056 (2011).

16 Chevalier, M., Gomez-Schiavon, M., Ng, A. & El-Samad, H. Design and analysis of a Proportional-Integral-Derivative controller with biological molecules. bioRxiv 303545; doi: https://doi.org/10.1101/303545 (2018).

17 Huang, H. H., Qian, Y. & Del Vecchio, D. A quasi-integral controller for adaptation of genetic modules to variable ribosome demand. Nat Commun 9, 5415, doi:10.1038/s41467-018-07899-z (2018).

18 Lillacci, G., Aoki, S. K., Schweingruber, D. & Khammash, M. A synthetic integral feedback controller for robust tunable regulation in bacteria. bioRxiv 170951; doi: https://doi.org/10.1101/170951 (2017).

19 Cardinale, S. & Arkin, A. P. Contextualizing context for synthetic biology - identifying causes of failure of synthetic biological systems. Biotechnol J 7, 856–866, doi:10.1002/biot.201200085 (2012).

20 Qian, Y. L. & Del Vecchio, D. Realizing ‘integral control’ in living cells: how to overcome leaky integration due to dilution? J R Soc Interface 15, doi:ARTN 2017090210.1098/rsif.2017.0902 (2018).

21 Rao, C. V., Wolf, D. M. & Arkin, A. P. Control, exploitation and tolerance of intracellular noise. Nature 420, 231–237, doi:10.1038/nature01258 (2002).

22 Wang, R. F. & Kushner, S. R. Construction of Versatile Low-Copy-Number Vectors for Cloning, Sequencing and Gene-Expression in Escherichia-Coli. Gene 100, 195–199 (1991).

23 Shin, J. & Noireaux, V. An E. coli Cell-Free Expression Toolbox: Application to Synthetic Gene Circuits and Artificial Cells. Acs Synthetic Biology 1, 29–41, doi:10.1021/sb200016s (2012).

24 Sun, Z. Z. et al. Protocols for Implementing an Escherichia coli Based TX-TL Cell-Free Expression System for Synthetic Biology. Jove-J Vis Exp, doi:ARTN e50762 10.3791/50762 (2013).

25 Maxwell, C. S., Jacobsen, T., Marshall, R., Noireaux, V. & Beisel, C. L. A detailed cell-free transcription-translation-based assay to decipher CRISPR protospacer-adjacent motifs. Methods 143, 48–57, doi:10.1016/j.ymeth.2018.02.016 (2018).

26 Agrawal, D. K. et al. Mathematical Modeling of RNA-Based Architectures for Closed Loop Control of Gene Expression. Acs Synthetic Biology 7, 1219–1228, doi:10.1021/acssynbio.8b00040 (2018).

27 Hu, C. Y., Varner, J. D. & Lucks, J. B. Generating Effective Models and Parameters for RNA Genetic Circuits. Acs Synthetic Biology 4, 914–926, doi:10.1021/acssynbio.5b00077 (2015).

28 Takahashi, M. K. et al. Characterizing and prototyping genetic networks with cell-free transcription-translation reactions. Methods 86, 60–72, doi:10.1016/j.ymeth.2015.05.020 (2015).

29 Westbrook, A. et al. Distinct timescales of RNA regulators enable the construction of a genetic pulse generator. bioRxiv 377572; doi: https://doi.org/10.1101/377572 (2018).

30 Frisk, A., Jyot, J., Arora, S. K. & Ramphal, R. Identification and functional characterization of flgM, a gene encoding the anti-sigma 28 factor in Pseudomonas aeruginosa. J Bacteriol 184, 1514–1521 (2002).

31 Slomovic, S., Pardee, K. & Collins, J. J. Synthetic biology devices for in vitro and in vivo diagnostics. Proc Natl Acad Sci U S A 112, 14429–14435. doi:10.1073/pnas.1508521112 (2015).

32 Bar-Ziv, R. Programmable on-chip DNA compartments as ‘artificial cells’. Eur Biophys J Biophy 46, S52–S52 (2017).

33 Caschera, F. & Noireaux, V. Compartmentalization of an all-E. coli Cell-Free Expression System for the Construction of a Minimal Cell. Artif Life 22, 185–195. doi:10.1162/ARTL_a_00198 (2016).

34 Majumder, S. et al. Cell-sized mechanosensitive and biosensing compartment programmed with DNA. Chem Commun 53, 7349–7352, doi:10.1039/c7cc03455e (2017).

35 Garamella, J., Marshall, R., Rustad, M. & Noireaux, V. The All E-coli TX-TL Toolbox 2.0: A Platform for Cell-Free Synthetic Biology. Acs Synthetic Biology 5, 344–355, doi:10.1021/acssynbio.5b00296 (2016).

36 Elowitz, M. B., Levine, A. J., Siggia, E. D. & Swain, P. S. Stochastic gene expression in a single cell. Science 297, 1183–1186, doi:DOI 10.1126/science.1070919 (2002).

37 Bleris, L. et al. Synthetic incoherent feedforward circuits show adaptation to the amount of their genetic template. Mol Syst Biol 7, doi:ARTN 51910.1038/msb.2011.49 (2011).

38 Segall-Shapiro, T. H., Sontag, E. D. & Voigt, C. A. Engineered promoters enable constant gene expression at any copy number in bacteria. Nat Biotechnol 36, 352, doi:10.1038/nbt.4111 (2018).

